# Extent and complexity of RNA processing in the development of honey bee queen and worker castes revealed by Nanopore direct RNA sequencing

**DOI:** 10.1101/2021.08.08.455492

**Authors:** Xu Jiang He, Andrew B. Barron, Liu Yang, Hu Chen, Yu Zhu He, Li Zhen Zhang, Qiang Huang, Zi Long Wang, Xiao Bo Wu, Wei Yu Yan, Zhi Jiang Zeng

**Affiliations:** Honeybee Research Institute, Jiangxi Agricultural University, Nanchang, Jiangxi 330045, P. R. of China; Jiangxi province Honeybee biology and beekeeping Nanchang, Jiangxi 330045, P. R. of China; Department of Biological Sciences, Macquarie University, North Ryde, NSW 2109, Australia; Wuhan Benagen Tech Solutions Company Limited, Wuhan, Hubei 430021, P. R. of China

**Keywords:** Honey bees, phenotypic plasticity, direct RNA sequencing, transcriptome complexity, isoform expression

## Abstract

The distinct honey bee (*Apis mellifera*) worker and queen castes have become a model for the study of genomic mechanisms of phenotypic plasticity. Prior studies have explored differences in gene expression and methylation during development of the two castes, but thus far no study has performed a genome-wide analysis of differences in RNA processing. To address this here we performed a Nanopore-based direct RNA sequencing with exceptionally long reads to compare the mRNA transcripts between honey bee queen and workers at three points during their larval development. We found thousands of significantly differentially expressed isoforms (DEIs) between queen and worker larvae. Most DEIs contained alternative splicing, and many of them contained at least two types of alternative splicing patterns, indicating complex RNA processing in honey bee caste differentiation. We found a negative correlation between poly(A) length and DEI expression, suggesting that poly(A) tails participate in the regulation of isoform expression. Hundreds of isoforms uniquely expressed in either queens or workers during their larval development, and isoforms were expressed at different points in queen and worker larval development demonstrating a dynamic relationship between isoform expression and developmental mechanisms. These findings show the full complexity of RNA processing and transcript expression in honey bee phenotypic plasticity.

## Introduction

Phenotypic plasticity - the capacity of divergent developmental pathways to yield more than one distinct phenotypes - is one of the most interesting aspects of development, but it remains poorly understood (Agrawal 2001; Borges 2005). The challenge is to understand how more than one developmental path can be organized and adaptively regulated by a single genome. Phenotypic plasticity is particularly dramatic in eusocial insects, and the honey bee (*Apis mellifera*) has become a valuable system for exploring this phenomenon. Honey bee queens and workers both develop from fertilized (diploid) eggs, but differences in nutrition in their larval diets determines their diverging developmental trajectories that result in different phenotypes (Asencot and Lensky 1984; Brouwers 1984; Maleszka 2008; Mao et al. 2015). The queen is the colony’s only fecund female with well-developed ovaries producing 1500-2000 eggs per day, whereas workers are infertile females that perform all the essential tasks in the colony such as brood care, foraging, defence, construction and cleaning (Winston 1991).

Prior foundational studies have shown how the process of phenotypic plasticity is epigenetically determined (Chen et al. 2012; Foret et al. 2012; Cameron et al. 2013; Guo et al. 2016; He et al. 2017; Wojciechowski et al. 2018). Studies have compared the transcriptomes between honey bee queens and workers using short-length RNA sequencing (RNA-Seq) (Chen et al. 2012; Cameron et al. 2013; He et al. 2017). Thousands of genes were differentially expressed between queens and workers during their development (Chen et al. 2012; Cameron et al. 2013; He et al. 2017). Consistently a few key signalling pathways have been implicated as important in queen/worker differentiation. These include target of rapamycin (TOR), the fork head box O (FoxO), Notch, Wnt, insulin/insulin-like signaling (IIS) pathways, mitogen-activated protein kinase (MAKP), Hippo, transforming growth factor beta (TGF-beta) (Patel et al. 2007; Mutti et al. 2011a; Duncan et al. 2016; Xiao et al. 2017; Yin et al. 2018; Wang et al. 2021); Key genes such as *Juvenile hormone esterase precursor (Jhe), ecdysone receptor* (*Ecr*), *Vitellogenin (Vg)*, *Hexamerin 70b* (*Hex70b*) *major royal jelly proteins* (*Mrjps*) also participate in the determination of queen-worker fate (Barchuk et al. 2007; Buttstedt et al. 2013; Mello et al. 2014). Moreover, epigeneic modifications such as DNA methylation, microRNA and alternative chromatin states have been shown to be involved in the regulation of honey bee queen-worker dimorphism (Foret et al. 2012; Guo et al. 2016; Wojciechowski et al. 2018). The latest evidence showed that m6A methylation on RNA also participates in the genomic regulation leading to queen-worker dimorphism (Wang et al. 2021).

These studies have gone a long way to establish the honey bee as a model for genomic analyses of phenotypic plasticity, but thus far, we know rather little about how RNA processing might be involved in the process. RNA processing determines the mRNA coding sequence, and can lead to different isoforms of a gene. Distinct isoforms can perform different biological functions, and these can be important for animal phenotypic plasticity (Marden 2008). Alternative transcripts produced from the same gene can differ in the position of the start site, the site of cleavage and polyadenylation, and the combination of exons spliced into the mature mRNA (Parker et al. 2020). Poly(A) tails are also very important for RNA processing. Length of the Poly(A) tail is correlated with translational efficiency and regulates transcript stability (Workman et al. 2019; Roach et al. 2020).

Alternative splicing (AS) of genes diversifies the transcriptome and increases protein coding capacity through the production of multiple distinct isoforms (Modrek and Lee 2001; Reddy et al. 2013). AS is involved into many biological processes in plants and animals, and particularly responses and adaptation to changing environments (Modrek and Lee 2001; Reddy et al. 2013). AS of pre-mRNA has been associated with phenotypic plasticity in insects such as the bumble bee *Bombus terrestris* and the pea aphid *Acyrthosiphon pisum* (Grantham and Brisson 2018; Price et al. 2018). Different alternative transcripts are also involved in honey bee caste differentiation, and alternative splicing plays an important role in queen-worker differentiation (Aamodt 2008; Mutti et al. 2011b). These studies have focused on the effects of AS on the regulation of gene expression. Thus far none have investigated how AS determines isoform expression genome wide in honey bees.

So far honey bee RNA-Seq studies have been based on short-length RNA sequencing, which is not an appropriate method to detect different isoforms or alternative transcripts of a gene. The new direct RNA sequencing technology (DRS) based on the Oxford Nanopore technology (ONT) directly sequences the native full-length mRNA molecules, which offers detailed information on the mRNA and RNA modifications (Garalde et al. 2018; Harel et al. 2019; Workman et al. 2019; Parker et al. 2020; Roach et al. 2020; Zhang et al. 2020). It avoids biases from reverse transcription or amplification and yields full-length, strand-special mRNA (Garalde et al. 2018; Harel et al. 2019; Zhang et al. 2020). This method allows a genome wide investigation of RNA processing [e.g. alternative splicing and poly(A) tails] and modifications (e.g. m6A).

Our objective in this study was to deploy this DRS technology to examine differences in RNA processing between the worker and queen honey bee castes. This allows us to investigate fundamental questions on caste-specific transcriptome patterns in honey bees: how isoforms are formed and expressed differentially in queens and workers, and the extent and complexity of the RNA processing in honey bee development.

We sampled honey bee queens and workers at different points in their larval development to assess the process of developmental canalization into the different phenotypes. Our results reveal extremely prevalent and highly complex differences in RNA processing between workers and queens. These differences changed across larval development showing a complex dynamical interaction between the epigenomic programs and the developmental pathways

## Results

### Quality of direct RNA sequencing data

Twelve libraries were generated and sequenced by ONT direct RNA sequencing. The average number of clean reads of each sample was 5,416,798 (Table 1). The average N50 read length was 1296 bp, and the average read quality was 10.63 (Table 1) indicating a high integrity of the sequenced RNAs. On average, 98.13% clean reads were successfully mapped to the *Apis mellifera* reference genome (*Amel HAv 3.1*) (Table 1). The Pearson correlation coefficients between two biological replicates of each group were all over 0.85 (Fig.S1) and the sample cluster tree also showed that two biological replicates of each group were clustered together (Fig.S2), reflecting a good repeatability of biological replicates.

**Table 1.** Overview of direct RNA sequencing

| Samples | 2Q-1 | 2Q-2 | 2W-1 | 2W-2 | 4Q-1 | 4Q-2 | 4W-1 | 4W-2 | 6Q-1 | 6Q-2 | 6W-1 | 6W-2 | Average |
| --- | --- | --- | --- | --- | --- | --- | --- | --- | --- | --- | --- | --- | --- |
| Clean read numbers | 7,268,687 | 5,793,325 | 6,113,875 | 6,093,102 | 6,067,308 | 6,945,178 | 6,207,113 | 3,807,886 | 5,039,594 | 2,057,391 | 5,910,366 | 3,697,756 | 5,416,798 |
| Average length | 1061.94 | 997.4 | 1001.55 | 1022.06 | 1007.76 | 998.23 | 967.9 | 1027.53 | 1036.56 | 1069.02 | 1027.87 | 976.65 | 1,016 |
| N50 length | 1356 | 1275 | 1285 | 1301 | 1295 | 1277 | 1242 | 1315 | 1338 | 1283 | 1306 | 1286 | 1,296 |
| Max read length | 23283 | 16804 | 16550 | 17946 | 29881 | 19966 | 16391 | 19519 | 17266 | 17829 | 13661 | 15496 | 18,716 |
| Average read quality | 10.71 | 10.61 | 10.75 | 10.71 | 10.26 | 10.53 | 10.66 | 10.62 | 10.77 | 10.68 | 10.66 | 10.64 | 10.63 |
| Alignment % identity | 98.80 | 98.35 | 98.42 | 98.46 | 98.23 | 98.63 | 98.19 | 98.38 | 96.85 | 98.40 | 98.71 | 96.19 | 98.13 |
| Uni mapped genes | 8,781 | 8,690 | 8,658 | 8,658 | 8,431 | 8,656 | 8,411 | 8,234 | 8,666 | 7,741 | 8,539 | 8,644 | 8,509 |
| Isoforms | 24,058 | 22,978 | 22,853 | 22,939 | 21,746 | 23,018 | 20,498 | 19,790 | 22,365 | 16,487 | 20,885 | 20,714 | 21,527 |

### Significantly differentially expressed isoforms (DEIs) and genes (DEGs) identified in queen-worker comparisons

Our results showed genome wide differentially expressed isoforms in honey bee caste differentiation. We identified 662, 1855 and 1042 DEIs in 2d, 4d and 6d queen-worker larval comparisons respectively, which were notably more than the DEGs (281, 1369 and 645 respectively, see Fig.1A, detailed information see Table S1-6). To compare the differences between DEIs and DEGs, we firstly mapped DEIs to reference genes (DEIGs) and showed that only 271 (34.72%), 977 (53.45%) and 410 (38.32%) genes overlapped between DEIGs and DEGs in 2d, 4d and 6d queen-worker comparisons respectively (Fig.1B,C and D). This shows a considerable difference between DEIs and DEGs in queen-worker differentiation.

**Fig.1A:**
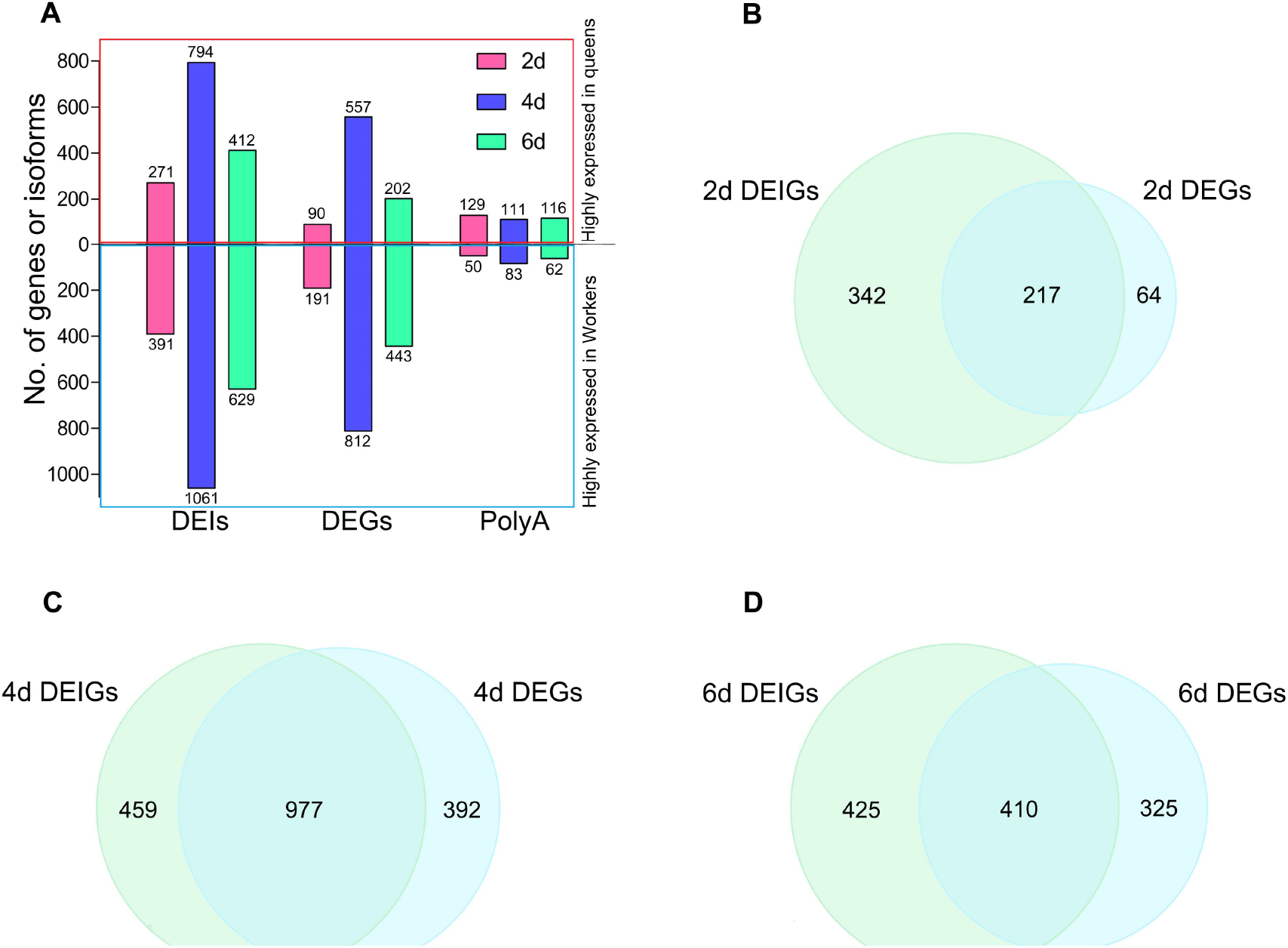
Numbers of differentially expressed isoforms (DEIs) and genes (DEGs), different poly(A)-length related isforms in three queen-worker larval comparisons (2d, 4d and 6d). The up bars are DEIs, DEGs and different poly(A) isoforms highly expressed in queens and the down bars are that highly expressed in workers. **B:** The venn diagram of DEIGs (DEI mapped genes) and DEGs from 2d queen-worker comparison. **C:** The venn diagram of DEIGs and DEGs from 4d queen-worker comparison. **D:** The venn diagram of DEIGs and DEGs from 6d queen-worker comparison.

We identified DEIs (24, 54, 48 in 2d, 4d and 6d comparisons respectively) that are involved in several key KEGG signaling pathways for honey bee caste differentiation, such as mTOR, Notch, FoxO, Wnt, IIS, Hippo, TGF-beta and MAPK (see Table S1-3). The number of DEGs enriched in these signal pathways were 7, 34, 33 in 2d, 4d and 6d respectively (Table S4-6). Other DEIs mapped to key genes such as *Jhe*, *Mrjps*, *Ecr*, *Vg* and *Hex70b* (see Table S1-3). The number of isoforms with significantly different poly(A) length in 2d, 4d and 6d comparisons showed a negative trend as DEIs and DEGs, with more in queens (Fig.1A, detailed information see Table S7-9) and less in workers.

DEIs and DEGs were enriched in five key KEGG signaling pathways (mTOR, Notch, FoxO, Wnt, IIS) which have been previous shown to be participating in of queen-worker differentiation. DEIs mapped to 35 enzymes in these five pathways, whereas DEG mapped onto just 17 enzymes (Fig.2A). In detail, 23 enzymes were uniquely mapped by DEIs whereas only 5 were uniquely mapped by DEGs. 12 enzymes mapped to both DEI and DEG (Fig.2A). We also provided detailed information about the expression of DEIs and DEGs enriched in these five key pathways (Fig.2B). Clearly, more isoforms were significantly differentially expressed in these five key pathways compared to DEGs, and the differentially expressed isoforms of each DEIG are presented in Fig.2B.

**Fig. 2A:**
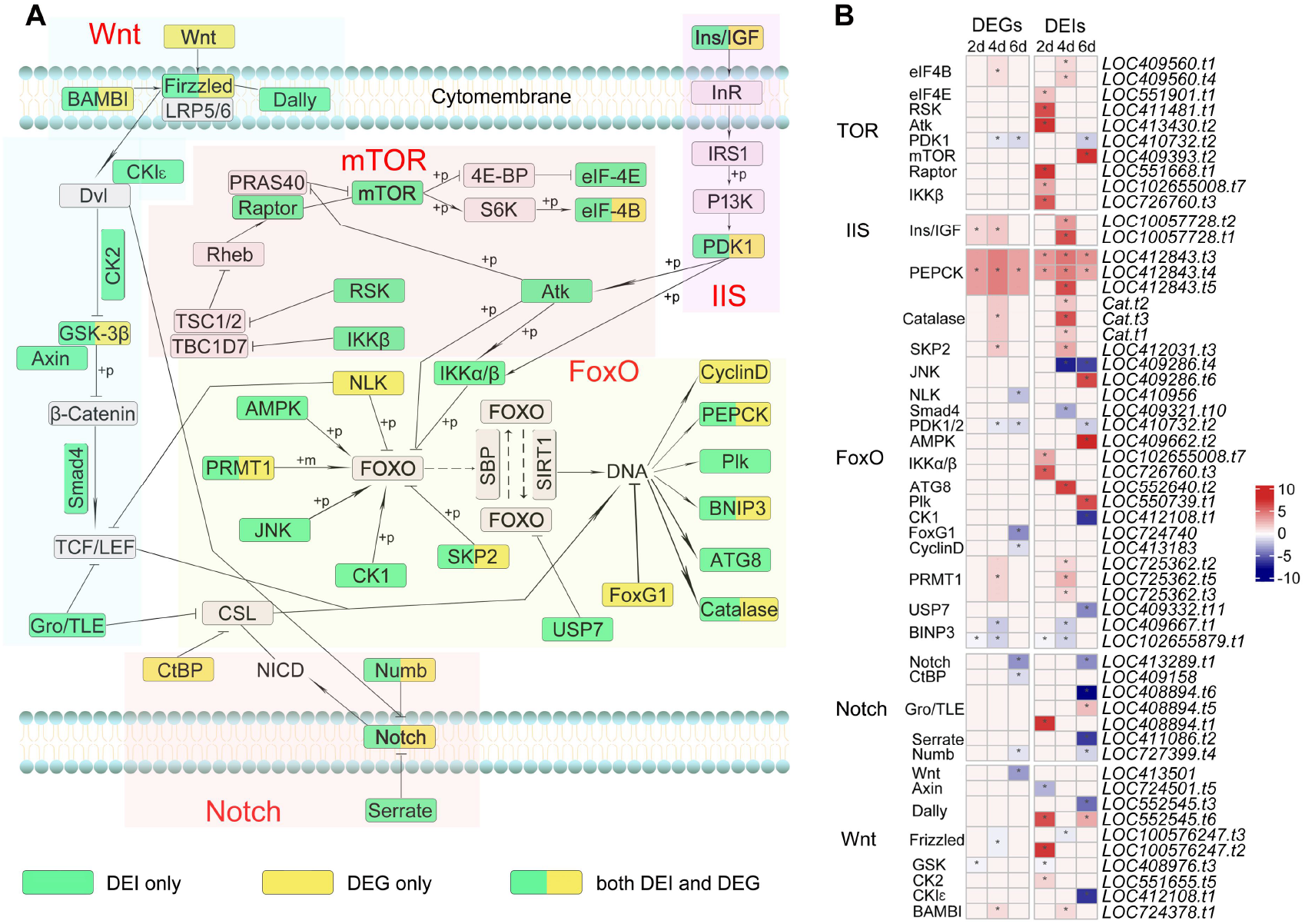
DEIs and DEGs enriched in key steps of five KEGG pathways. Five KEGG pathways were shown with transparent boxes and the names of these pathways were red marked. Green boxes inside of transparent boxes are DEI enriched enzymes; yellow boxes are DEG enriched enzymes; green-yellow mixed boxes are both DEI and DEG enriched enzymes; blank boxes are non DEI and DEG enriched enzymes. **B:** Expression of DEIs and DEGs in the five key pathways. The left are pathway and enzyme names, and the right are isoform expression and names. Expression of isoforms and genes are presented with their log2TPM values and shown with color scales. DEIs or DEGs are marked with “*” in the middle of the boxes.

These findings suggest that caste-specific isoform expression provided more accurate and detailed information of the real transcriptome differences between honey bee queen and worker castes than measurements of differences in gene expression.

### Uniquely expressed isoforms between queens and workers

We identified 187, 357 and 364 uniquely expressed isoforms in queen or worker larvae at 2d, 4d and 6d age, with more occurring in queens than workers (Fig. 3A). The number of uniquely expressed isoforms in queens reached a maximum at the 4d stage whereas in workers it reached a maximum in the 6d sample. This could suggest that the queen developmental pathway diverges more quickly and at an early larval stage while the developmental pathway of workers diverges slightly later.

**Fig. 3A:**
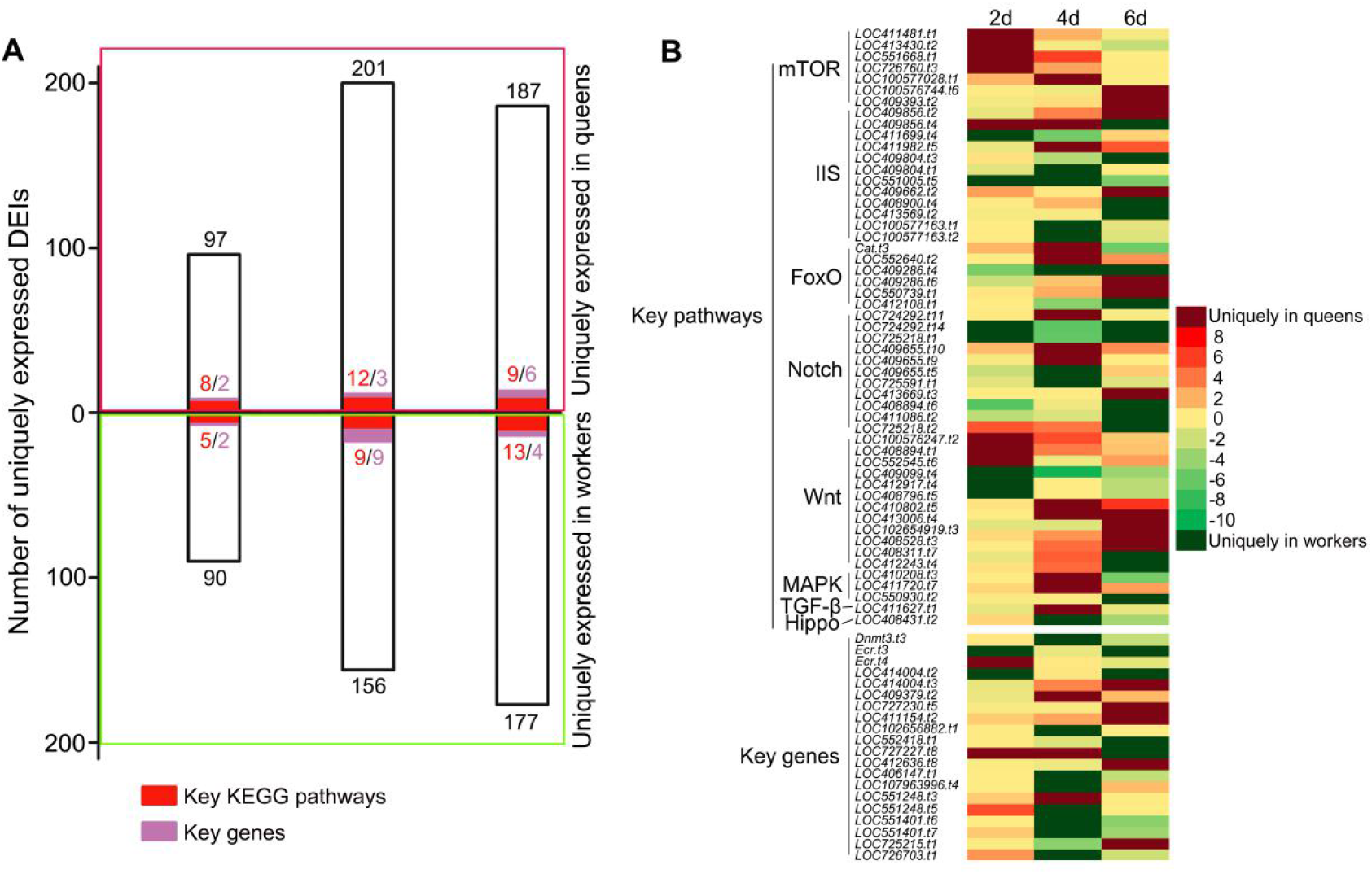
Numbers of uniquely expressed isoforms in queen larvae (up bars) or worker larvae (down bars) at 2d, 4d and 6d stages. The red bars are numbers of uniquely expressed isoforms enriched in eight key KEGG signaling pathways (mTOR, IIS, FoxO, Notch, Wnt, MAPK, Hippo and TGF-beta). The purple bars are numbers of uniquely expressed isoforms that are key genes reported in previous studies (details see Table S10-12). **B:** The heat map of expression of uniquely expressed isoforms which are shown in red bars and green bars in A. The red color blocks in heat map are uniquely expressed isoforms in queen larvae at 2d, 4d and 6d age, whereas the dark green blocks are uniquely expressed isoforms in worker larvae. Other color scales are the log2fold change values of isoform expression between queen and worker larvae at 2d, 4d and 6d stages.

Uniquely expressed isoforms were enriched into eight key KEGG signaling pathways such as mTOR and IIS for queen-worker differentiation as well as some key genes such as *DNA methyltransferase 3 (Dnmt3)* and *<u>Ecr</u>* (Fig.3, details see Table S10-12). This indicated that there are isoforms which are likely to be involved in queen-worker differentiation that are uniquely expressed in queens or workers during their development.

### DEIs and alternative splicing

We identified 21,574 isoforms in each sample on average, which was three times more than the number of protein-coding genes (on average 8509) (Table 1). This suggested that the honey bee genome produces various different isoforms from one gene. We therefore detected AS events in the DEIs. The majority of DEIs (73.26%, 67.98% and 63.53% of 2d, 4d and 6d DEIs) contained AS events, and most of them (71.13%, 66.16% and 65.56% in 2d, 4d and 6d) had at least two types of AS in a single isoform. A combination of RI, A5 and A3 in a single DEI was the most common type (Fig.4). This indicates that the formation of transcripts in honey bee development is more complex than previous known.

**Fig. 4.**
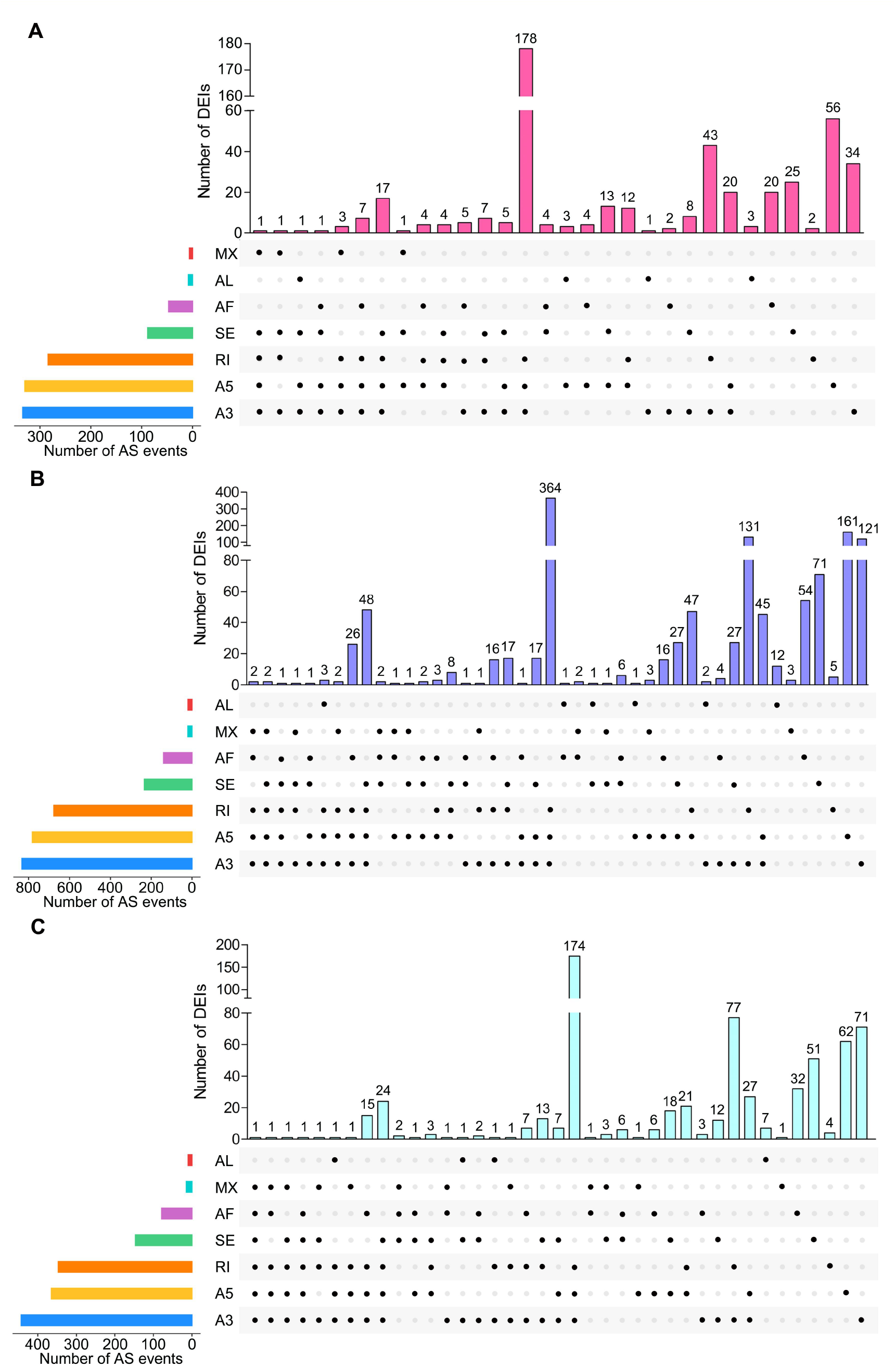
The alternative splicing patterns (AS) in DEIs of three comparisons. **A:** DEIs of 2d comparison containing AS events. The colorful bars in the left bottom diagram are the different AS patterns and the number of AS events in DEIs. Seven different AS patterns are 3’splice site (A3), 5’splice site (A5), First exon (AF), Last exon (AL), Retained intron (RI), Skipping exon (SE) and Mutually exclusive exon (MX). The pink bars in the top of right diagram are the number of DEIs. The black spots in a column mean one single isoform containing these different AS patterns. Same in B and C. **B:** DEIs of 4d comparison containing AS events. **C:** DEIs of 6d comparison containing AS events.

Of these DEIs, 9.82% (65), 5.71% (106), 9.50% (99) of DEIs from the 2d, 4d and 6d comparisons contained significantly different AS events when comparing workers and queens (Fig. S3, Table S13-15). Similarly, many of them (46.15%, 42.45% and 45.45% in 2d, 4d and 6d comparisons respectively) contained at least two types of significantly different AS events (Fig. S4, Table S13-15). Consequently, a part of DEIs were significantly differentially spliced.

### Correlation between poly(A) length and expression of DEIs and DEGs

We showed a negative correlation between isoform expression and their poly(A) tail length (Fig. S5). Expression of DEIs (Fig.5) and DEGs (Fig.S6) was also negatively correlated with their poly(A) lengths. It is believed that poly(A) tails are involved in the regulation of isoform expression.

**Fig. 5:**
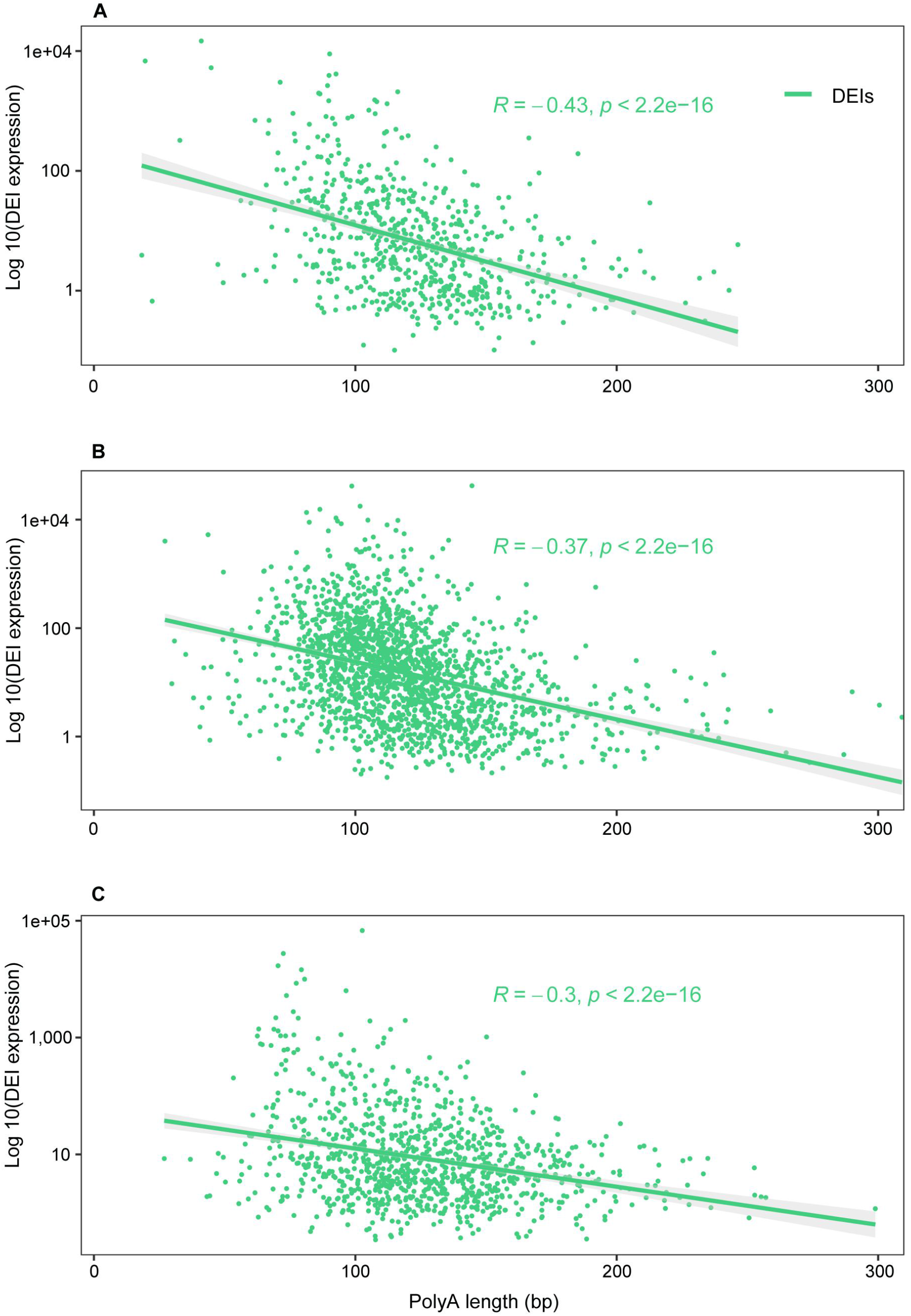
Correlation between poly(A) lengths and DEI expression of 2d queen-worker comparison. Expression of each DEI was the log10 TPM value. Same in B and C. **B:** Correlation between poly(A) lengths and DEIs of 4d comparison. **C:** Correlation between poly(A) lengths and DEIs of 6d comparison.

### Correlation of isoform expression and poly(A) lengths for two key genes

Two key genes which participate in queen-worker differentiation are *Jhe* and *Ecr*. These were selected as examples to show the correlation between isoform expression and poly(A) tails. *Jhe* had only one isoform, and was negatively correlated with its poly(A) lengths (Fig. 6A). The *Ecr* gene had 4 isoforms. We selected the most differentially expressed isoform (*Ecr.t4*) to measure the correlation between its expression and poly(A) length. Similarly, the expression of *Ecr.t4* isoform was also negatively correlated with their poly(A) lengths (Fig. 6B).

**Fig. 6A:**
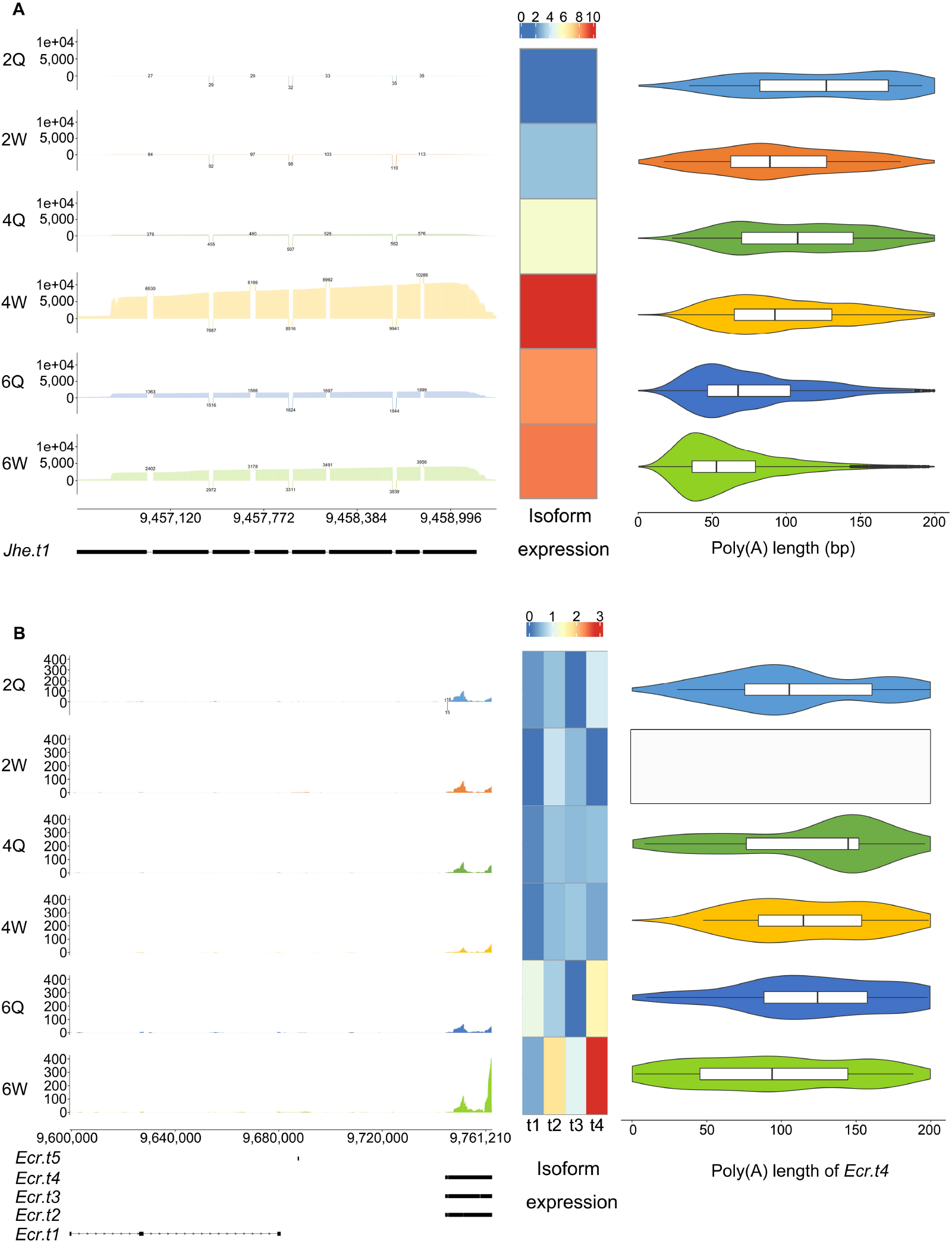
Expression and related poly(A) lengths of *Jhe* isoform. The left is the exon coverage of clean reads of *Jhe* gene of 6 larval groups. The *Jhe* has only one isoform (*Jhe.t1*) and its structure is present (see its exons below marked with black color). The middle heat map is the isoform expression of *Jhe.t1* (log2TPM values) in 6 groups and presented with color scales. The right is the poly(A) length distribution of *Jhe.t1* in 6 groups. The mean poly(A) length is presented in the middle of each violinplot. **B:** Expression and related poly(A) lengths of *Ecr* isoforms. The *Ecr* gene has 4 isoforms and their expression is presented using heat map as above. The poly(A) length distribution of *Ecr.t4* in 6 groups is presented on the right side, it was the most significantly differentially expressed isoform of gene *Ecr*. The poly(A) of *Ecr.t4* in 2dW was presented with a blank box which means few clean data of poly(A) was detected, since 2dW did not express *Ecr.t4*.

## Discussion

RNA processing act as a major driver of animal phenotypic plasticity (Gingeras 2007), but the complexity and characteristics of RNA processing underlying this phenotypic variability is not well understood. The present study has shown the extent of RNA processing in honey bee queen-worker differentiation. Differences in isoform expression revealed the true extent of the differences in the transcriptome between queens and workers during their development, providing more detailed and accurate information on mRNA expression compared to measures of gene expression. We also showed the complexity of alternative splicing patterns in queen-worker differentiation. Most DEIs were formed by more than one type of alternative splicing. Poly(A) length negatively correlated with DEIs expression demonstrating another part of the genomic mechanisms of honey bee caste differentiation.

Thousands of DEIs were identified between queens and workers, which were notably more than DEGs (Fig.1A). Many genes were not differentially expressed, but different isoforms from these genes were significantly differentially expressed (Fig. 1B-D, Table S1-3), demonstrating how DRS gives a very different perception of the transcriptome differences between developing worker and queen honey bees. DEIs enriched into more enzymes of five key KEGG signaling pathways such as mTOR and IIS than DEGs (Fig 2). The mTOR signaling pathway is the central component of a conserved eukaryotic signaling pathway, regulating cell and organismal growth in response to nutrient status and has been shown to be important for determining the queen-worker developmental paths (Colombani et al. 2003; Oldham and Hafen 2003; Patel et al. 2007). The IIS and Notch signaling pathway have been shown to be involved in the development of honey bee ovaries (Mutti et al. 2011a; Duncan et al. 2016). More DEIs were identified in these key pathways than DEGs, indicating that DEIs could provide more nuanced differences of mRNA expression between honey bee queens and workers during their development. Therefore, compared to DEGs that conceals a more nuanced picture of transcription differences, DEIs show the true nature of transcriptome differences between queens and workers. Consequently, DEIs identified in the present study revealed a more informative knowledge and precise difference in the transcriptome expression of honey bee caste differentiation compared to previous RNA-Seq results (Chen et al. 2012; Cameron et al. 2013; He et al. 2017; Yin et al. 2018).

Hundreds of uniquely expressed isoforms were identified in queens or workers. Many uniquely expressed isoforms were enriched in mTOR, IIS, Notch, FoxO etc eight key KEGG signaling pathways (Fig.3). Some uniquely expressed isoforms mapped to key genes such as DNA methyltransferase 3 (*Dnmt3*) and *Ecr* genes (Fig.3). *Dnmt3* is a key driver of epigenetic global reprogramming controlling honey bee queen-worker fate (Kucharski et al. 2008). Here we showed one isoform of *Dnmt3* (*Dnmt3.t3*) was uniquely expressed in 4d worker larvae. It is possible that this isoform of *Dnmt3* in worker larvae modifies the DNA methylation status and helps to canalize the worker developmental pathway. For the *Ecr* gene, 2d worker uniquely expressed *Ecr.t3* whereas 2d queen larvae uniquely expressed *Ecr. t4*. Different isoforms from the same gene were expressed in queens and workers, revealing the complexity of of RNA processing in the honey bee queen-worker differentiation. We have documented many genes where, isoform expression was not consistently biased toward either queens or workers throughout all stages of larval development. In some cases, an isoform was uniquely expressed in either queens or workers in one larval stage, but at other developmental stages the isoform occurred in both workers and queens (Fig. 3B). Isoforms like LOC409856.t4 and LOC727227.t8 uniquely expressed in queens at 2d and 4d stages but oppositely expressed in workers at 6d stage (Fig. 3B). This illustrates the dynamic relationship between RNA processing and the developmental process. A simple model might imagine different isoforms of a gene specific to either the queen or worker developmental pathway. Instead we see isoforms featuring in both pathways, but at different stages.

The number of uniquely expressed isoforms also showed a different trend in queens and workers. It reached a maximum in queens at the 4d larval stage but it peaked later in workers (Fig. 3A). These different trends correlate with the known faster development of queens (Winston 1991) and suggest the queens developmental pathway differentiates at an earlier larvae stage than the worker developmental pathway. Queen larvae at early stage receive significantly more food than worker larvae, and the nutrient contents in royal jelly also differ from worker jelly in terms of sugar, vitamins, proteins, acids, microRNA (Asencot and Lensky 1984; Brouwers 1984; Maleszka 2008; Mao et al. 2015). In Fig. 3B, most isoforms enriched in mTOR signaling pathway which is a signaling pathway in response to nutrient status (Colombani et al. 2003; Oldham and Hafen 2003) were uniquely expressed in queens at 2d stage and its number decreased at 4d and 6d stages, which supports our speculation.

We detected on average 21,574 isoforms from 8509 genes in each sample (Table 1). This is consistent with a previous study that 60.3% of honey bee multi-exon genes are alternatively spliced (Li et al. 2013). The majority of DEIs, contained AS events and over 65% of them had at least two types of AS patterns in a single isoform (Fig. 4). Some DEIs contained more than five different AS types (Fig. 4). A few DEIs were significantly differently spliced between queens and workers by multiple AS patterns (Fig. S3, table S13-15). It means several distinct AS types together contributed to the construction of a single isoform (Fig.4), revealing an extremely complex splicing system in honeybee caste differentiation.

Poly(A) tails are a regulator of translation and transcript stability (Lim et al. 2016; Tudek et al. 2018; Woo et al. 2018). Poly(A) tail length is not stable but dynamic and condition-dependent (Lim et al. 2016; Woo et al. 2018). This study showed that poly(A) tail lengths were strongly and negatively correlated with expression of isoforms in honey bees (Fig. 5). The negative correlation between isoform expression and poly(A) length has been shown in the regulation of *Caenorhabditis elegans* development (Roach et al. 2020). Here we showed that poly(A) tails also participate in the regulatiion of isoform expression, and therefore contributes to complexity of RNA processing in honeybee queen-worker differentiation.

In summary, RNA processing plays an important role in shaping honey bee phenotypic plasticity, but investigations of gene expression by short-length RNA-Seq fail to reveal the full complexities of RNA processing (Chen et al. 2012; Cameron et al. 2013; He et al. 2017; Yin et al. 2018). By using DRS, this study has shown the extent of differential isoform expression between workers and queens, complexities of transcript splicing, and polyadenylation. It provides a more detailed understanding of the molecular mechanisms underlying the divergence of the queen and worker developmental paths. It also contributes to our understanding of the extent and complexity of RNA processing in animal phenotypic plasticity.

## Material and methods

### Insects

Three healthy honey bee colonies (*Apis mellifera*) each with a mated queen were used for this study. Each colony had ten frames with approximately 35,000 bees. These colonies were maintained at the Honeybee Research Institute, Jiangxi Agricultural University, Nanchang, China, according to the standard beekeeping techniques.

The mother queen was caged onto a plastic frame (Pan et al. 2013) to lay eggs into worker cells for 6 hours. Afterwards, half of these eggs were removed into commercial plastic queen cells to rear queens before hatching. Queen and worker larvae at 48 hrs (2 day), 96hrs (4 day) and 144 hrs (6 day) after hatching were harvested and were flash-frozen in liquid nitrogen. In the DRS experiment, for the 2dQL or 2dWL, larvae were very small, therefore we collected 20 2d queen larvae and mixed them into one sample. The 4d and 6d larvae were large enough, and one sample needed only one larva. Each group had two replicates from two colonies, therefore in total 12 samples were collected. Larval samples for short-length RNA-Seq were sampled the same as DRS, but with twice the number of larvae in each sample. We had three replicates for each of these 6 groups: 18 samples in total. Within a colony queens and workers developed from a same mother queen and eggs were laid at the same time. All sampled eggs were reared in a same colony. We compared queen and worker larvae sampled at the same time point: 2dQL vs 2d WL, 4dQLvs 4dWL, and 6dQL vs 6dWL.

### RNA preparation

Total RNA of 2^nd^, 4^th^ and 6^th^ instars of queens and workers as extracted in accordance with the standard protocol of the of the Total RNA Kit I (R6834, Omega Bio-tek), respectively. Each sample contained 30 μg total RNA for DRS. RNA was cleaned and concentrated by NEBNext Poly(A) mRNA Magnetic Isolation Module (E7490S) according to the manufacturer’s instructions. From each sample we extracted 1 μg of total RNA for the RNA integrity and concentration measurement using Nanodrop (Thermo Fisher Scientific).

### Direct RNA sequencing

Prepared RNA of each sample was used for a DRS library preparation using the Oxford Nanopore DRS protocol (SQK-RNA002, Oxford Nanopore Technologies). For reversed connector connection, 9 μL prepared RNA, 3μL NEBNext Quick Ligation Reaction Buffer(NEB), 1 μL RT Adapter (RTA)(SQK-RNA002) and 2 μL T4 DNA Ligase (NEB) were mixed together and incubated under 25 °C for 10 min. Afterwards, 8 μL 5x first-strand buffer (NEB), 2 μL 10 mM dNTPs (NEB), 9 μL Nuclease-free water, 4 μL 0.1 M DTT (Thermo Fisher) and 2 μL SuperScript III Reverse Transcriptase (Thermo Fisher Scientific, 18080044) were added into the above 15 μL reaction system and incubated under 50 °C for 50 min then 70 °C for 10min. Reverse-transcribed mRNA was purified with 1.8*Agencourt RNAClean XP beads and washed with 23ul Nuclease-free water, and subsequently sequencing adapters were added using 8 μL NEBNext Quick Ligation Reaction Buffer, 6 μL RNA Adapter (RMX) and 3 μL T4 DNA Ligase. The mix was purified and washed again as above, and 75 μL RRB (SQK-RNA002) were added with 35 μL Nuclease-free water. This final reaction system was loaded into Nanopore R9.4 sequencing micro-array and sequenced for 48-72 hrs using PromethION sequencer (Oxford Nanopore Technologies). This was performed by Wuhan Benagen Tech Solutions Company Limited.

### Preprocessing and alignments

To assess the read quality, raw reads were base called using Guppy software (version 3.2.6) (Fesenko and Knyazev 2020) and adapter sequences were trimmed by Guppy as well. Low quality reads (Q-value < 7) and short-length reads (<50 bp) were filtered by Nanofilt (version 2.7.1) (Coster et al. 2018). The remaining clean reads were corrected using the Fclmr2 (version 0.1.2) with our short-length RNA-Seq data (Wang et al. 2018). Afterwards, clean reads from each library were aligned to the genome of honey bees *Apis mellifera* (*Amel* HAv3.1) using Minimap2 (version 2.17-r941) (Li 2018). The alignment ratio of clean reads to the honey bee reference genes were calculated using Samtools (version 1.10) (Li et al. 2009) and showed in Table 1. After clearing redundant sequences using Flair collapse (version 1.5.0) (Tang et al. 2020) and Stringtie (version 2.1.4) (Pertea et al. 2015), the clean reads from each library were merged by Gffcompare (version 0.12.1) (Pertea and Pertea 2020) to obtain the final isoforms.

### DEG and DEI calculation

Isoform expression was annotated to the honey bee reference genes and quantified using the selective-alignment-base model in Salmon software (version 1.4.0) (Patro et al. 2017). Gene expression was measured according to standard methods as in He et al (He et al. 2017) using DRS raw data rather than RNA-Seq data. Significantly differentially expressed genes (DEGs) and isoforms (DEIs) between queen and worker larvae were identified as those with p-values <0.05 and log2 Fold change values > 1.5 using DESeq2 (version 1.26.0) (Love et al. 2014).To compare the expression level of genes/isoforms, we computed TPM values (transcripts per kilobase of exon model per million mapped reads) based on reads counts as expression. To compare the DEGs and DEIs, all DEIs were referenced to the genes to obtain Differentially Expressed Isoform of Genes (DEIGs). Sequences of the DEGs and DEIGs were blasted to the Swiss-Prot database, non-redundant protein (Nr) and non-redundant nucleotide sequence (Nt) database with a cut-off E-value of 10^−5^.

### KEGG pathway enrichment analysis

Five KEGG pathways [TOR, FoxO, Notch, Wnt and IIS] which have been repeatedly associated with honey bee queen-worker differentiation (Patel et al. 2007; Mutti et al. 2011a; Duncan et al. 2016; Xiao et al. 2017; Yin et al. 2018; Wang et al. 2021) were selected and combined into a pathway net according to KEGG pathway database and a previous study (Wang et al. 2021). The statistical enrichment of DEIGs and DEGs in these KEGG pathways were analyzed using a hypergeometric test (Q-value < 0.05) using the KOBAS 2.0 software (Xie et al. 2011). Results were shown in Fig. 3A. Expression of DEIs and genes in these five pathways were also presented in Fig.3B.

### Alternative splicing identification

The AS events of each comparison were analyzed using SUPPA2 (version 2.3) (Trincado et al. 2018). AS events were divided into seven patterns including 3’splice site (A3), 5’splice site (A5), First exon (AF), Last exon (AL), Retained intron (RI), Skipping exon (SE) and Mutually exclusive exon (MX). The PSI values of each AS events in DEIs were calculated after filtering percent spliced in index (PSI) values with ‘not available’ (NA) or ‘0’ (Trincado et al. 2018). Significant differences in AS events were identified as its dPSI-value >0 and p-value <0.05.

### Poly(A) length estimation and correlation with expression of genes and isoforms

Poly(A) lengths of each read were calculated by the Nanopolish software (version 0.13.2) (Coster et al. 2018). The 3’ poly(A) tail lengths of each read were directly identified using the ionic current signals (Workman et al. 2019). A previous study showed that poly(A) lengths were negatively correlated with isoform expression, therefore, the median of poly(A) length of each isoform was used for correlation and regression analysis with their TPM values (results were shown in Fig.S5) in R package (version 4.0.3). We also evaluated the correlation between poly(A) tail lengths and DEIs or DEGs.

For this we used median of poly(A) length and TPM or log2Fold change values of DEIs or DEGs for correlation and regression analysis as above. Two key genes (*Jhe* and *Ecr)* for the determination of queen-worker differentiation were selected as an example to show the correlation between expression of isoforms and poly(A) lengths. Results were shown in Fig. 5.

### Short-length RNA sequencing

Total RNA of each sample was extracted using TRIzol reagent (Tiangen, Beijing). The concentration and integrity of RNA were assessed using Qubit 2.0 Fluorometer and Agilent 2100 bioanalyzer. Each group of queen and worker larvae had 3 biological replicates.

Total mRNA of each sample was enriched using oligo (dT) magnetic beads. First-strand cDNA was synthesized by random hexamers, and the second-strand cDNA was synthesized in DNA polymerase I system using dNTPs and RNaseH. Afterwards, cDNA was subjected to end repair and performed with poly(A) tail and ligation sequencing adapter. 200-350 bp cDNA were selected by AMPure XP beads, and then were PCR amplified and purified by AMPure XP beads. Library quality was evaluated using Agilent 2100 bioanalyzer and qRT-PCR. Totally 18 libraries were sequenced by an Illumina NovaSeq 6000 platform. This was performed by Wuhan Benagen Tech Solutions Company Limited also.

## Data access

All data are available from the corresponding author upon request. The DRS sequence data for all 12 libraries and the fastq sequences of RNA-Seq of 18 libraries have been deposited in the NCBI database under the accession: NCBI Bioproject: PRJNA748829.

## Acknowledgement

We thank Mr Yi Cheng Yang and Mrs Yan Niu for help on data analysis and Mrs Xin Zhang for help on project management. This work was supported by the National Natural Science Foundation of China (31702193, 31872432), Jiangxi provincial academic and technical leader project (20204BCJL23041) and the Earmarked Fund for China Agriculture Research System (CARS-44-KXJ15).

## Authors’ contribution

Z.J.Z. and X.J.H. designed research; X.J.H. performed the research; X.J.H., A.B.B. provided guidance for data; X.J.H., L.Y., and H.C. analyzed data; Y.Z.H., L.Z.Z., Q.H., Z.L.W., X.B.W., and W.Y.Y. assisted in the experiments. X.J.H., A.B.B. and Z.J.Z. wrote the paper.

## Supplementary

Fig. S1 The pearson correlation coefficient of 2d, 4d and 6d queen (QL) and worker larvae (WL). Each larval group had two biological replicates (A and B).

Fig. S2 The sample cluster tree of 12 samples. Isoform expression of each sample was compared and its correlation was presented using color scales.

Fig. S3 DEIs containing significantly different AS events in 2d, 4d and 6d queen-worker comparisons. Blue color presents the proportion of DEIs containing AS events, and red color presents DEIs containing significantly different AS events.

Fig. S4 The significantly differential alternative splicing patterns in DEIs of three comparisons (similar to Fig.4). **A:** DEIs of 2d comparison containing significantly differential AS events. The colorful bars in the left bottom diagram are the different AS patterns and the number of AS events in DEIs. Seven different AS patterns are 3’splice site (A3), 5’splice site (A5), First exon (AF), Last exon (AL), Retained intron (RI), Skipping exon (SE) and Mutually exclusive exon (MX). The pink bars in the top of right diagram are the number of DEIs. The black spots in a column mean one single isoform containing these different AS patterns. Same in B and C. **B:** DEIs of 4d comparison containing significantly differential AS events. **C:** DEIs of 6d comparison containing significantly differential AS events.

Fig. S5 Correlation between poly(A) lengths and isoform expression of 12 samples. Expression of each isoform was the log10 TPM value.

Fig. S6 Correlation between poly(A) lengths and DEG expression of three queen-worker comparisons. Up: 2d comparison, middle: 4d comparison, down: 6d comparison. Expression of each DEG was the log10 TPM value.

Table S1 DEIs between 2d queen and worker larvae

Table S2 DEIs between 4d queen and worker larvae

Table S3 DEIs between 6d queen and worker larvae

Table S4 DEGs between 2d queen and worker larvae

Table S5 DEGs between 4d queen and worker larvae

Table S6 DEGs between 6d queen and worker larvae

Table S7 Different poly(A) related genes between 2d queen and worker larvae

Table S8 Different poly(A) related genes between 4d queen and worker larvae

Table S9 Different poly(A) related genes between 6d queen and worker larvae

Table S10 Uniquely expressed isoforms in 2d queen or worker larvae

Table S11 Uniquely expressed isoforms in 4d queen or worker larvae

Table S12 Uniquely expressed isoforms in 6d queen or worker larvae

Table S13 DEIs of 2d comparison containing significantly different AS events

Table S14 DEIs of 4d comparison containing significantly different AS events

Table S15 DEIs of 6d comparison containing significantly different AS events

