## Supplementary figures and images for "Extent and complexity of RNA processing in the development of honey bee queen and worker castes revealed by Nanopore direct RNA sequencing"

### Supplemenntary Figure1-6

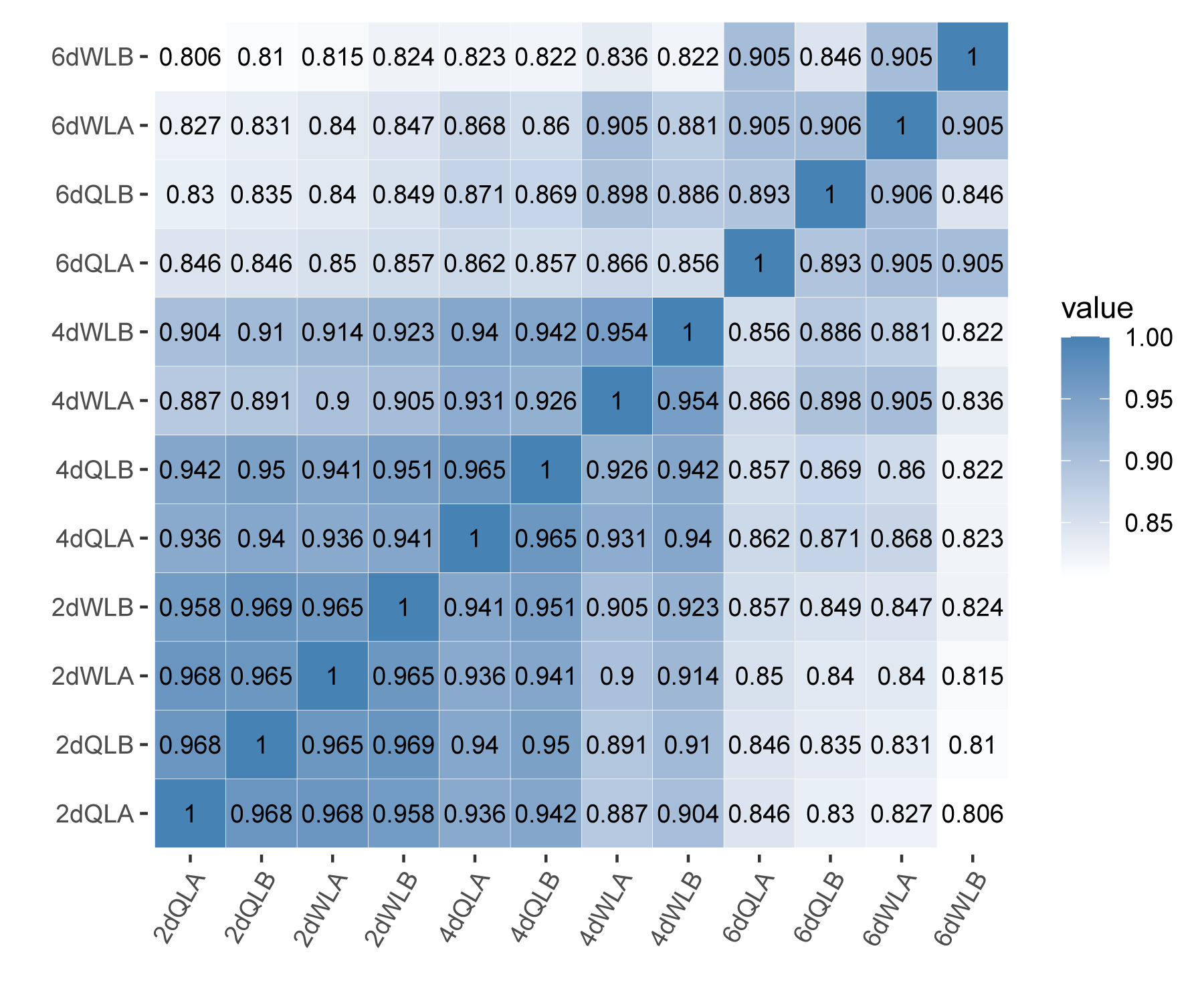

Fig.S1

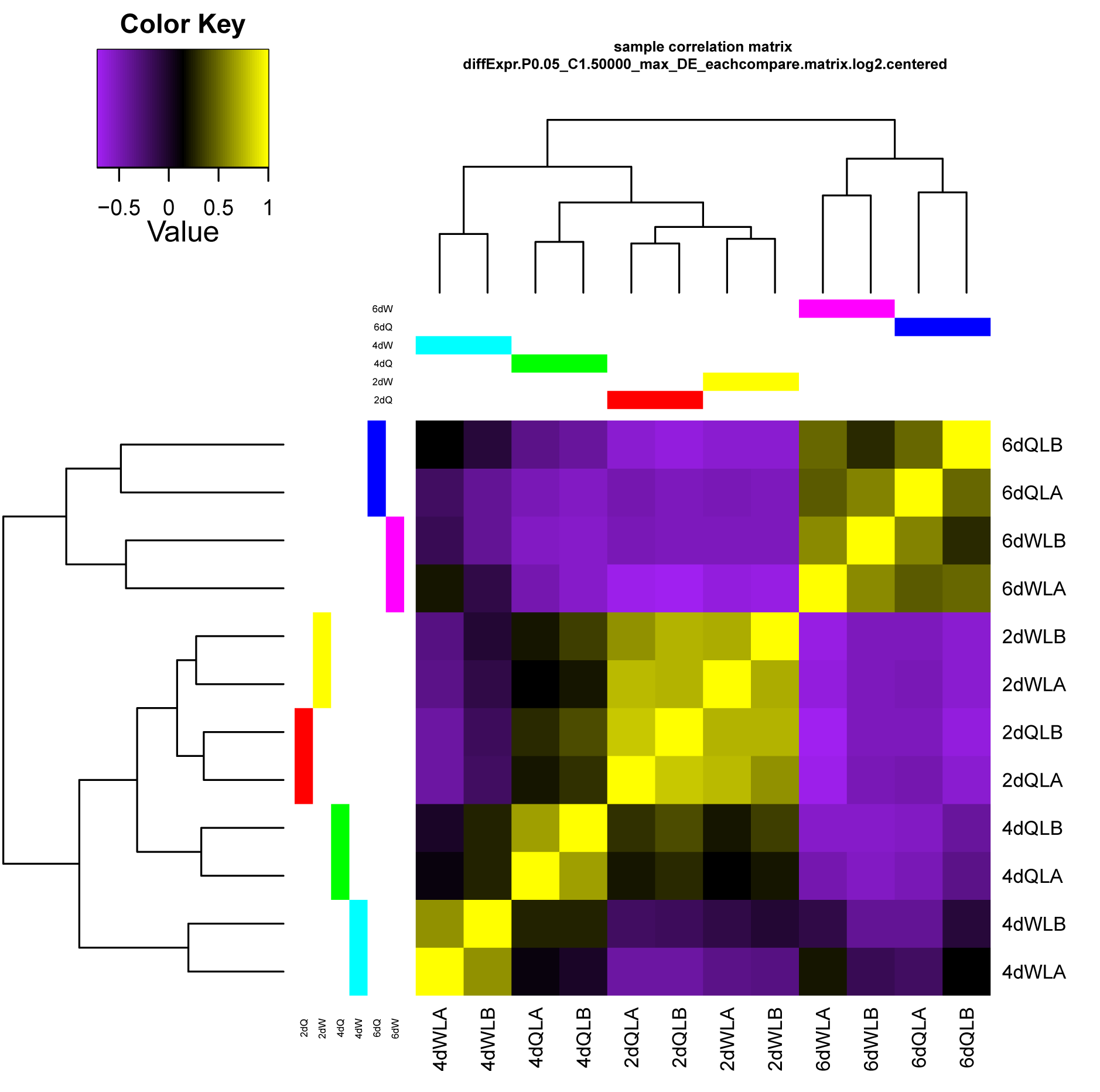

Fig.S2

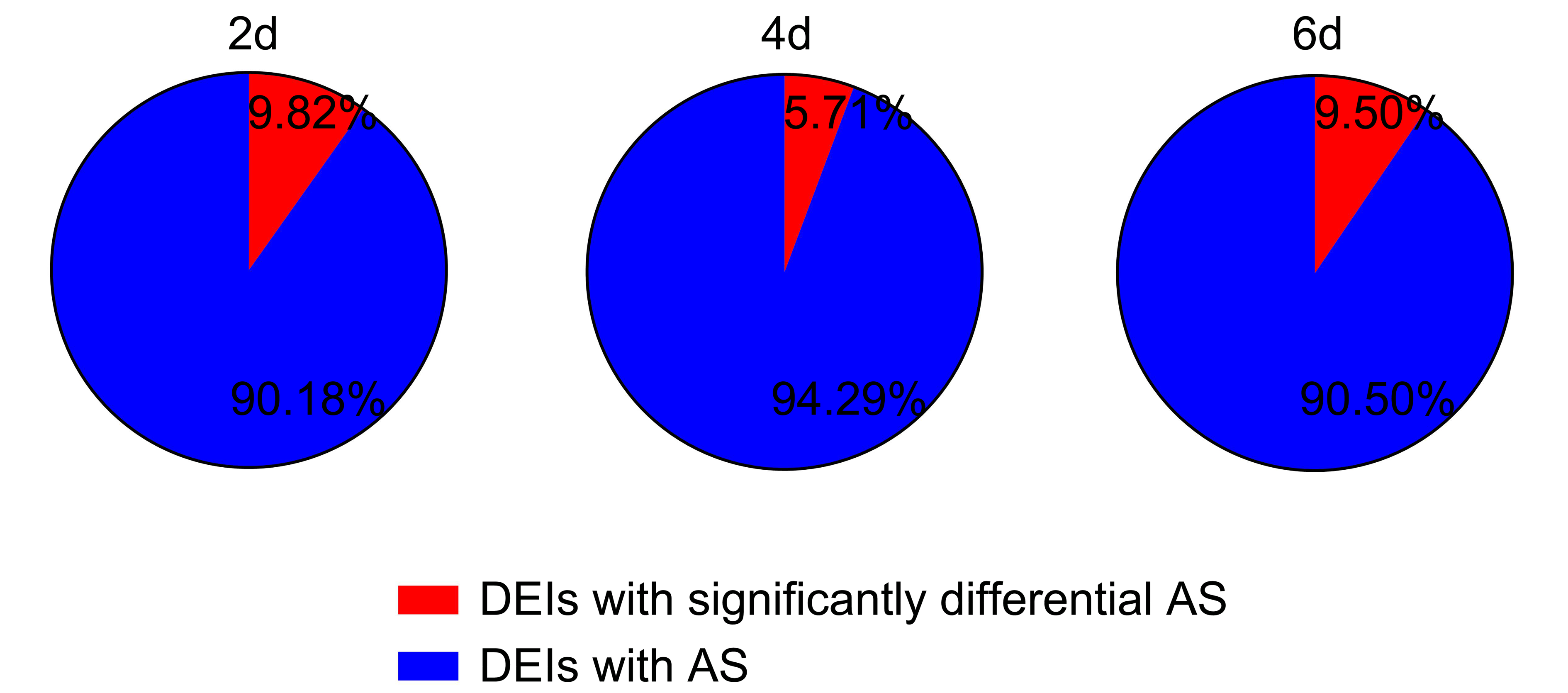

Fig.S3

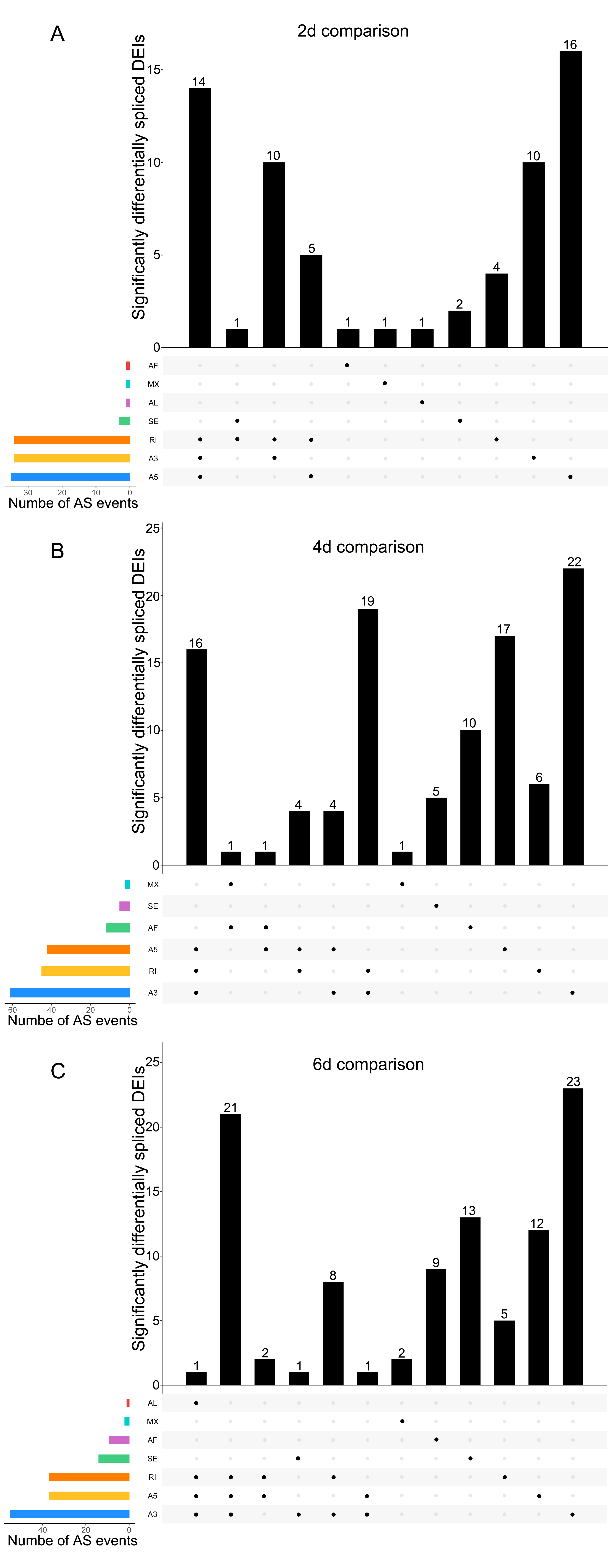

Fig.S4

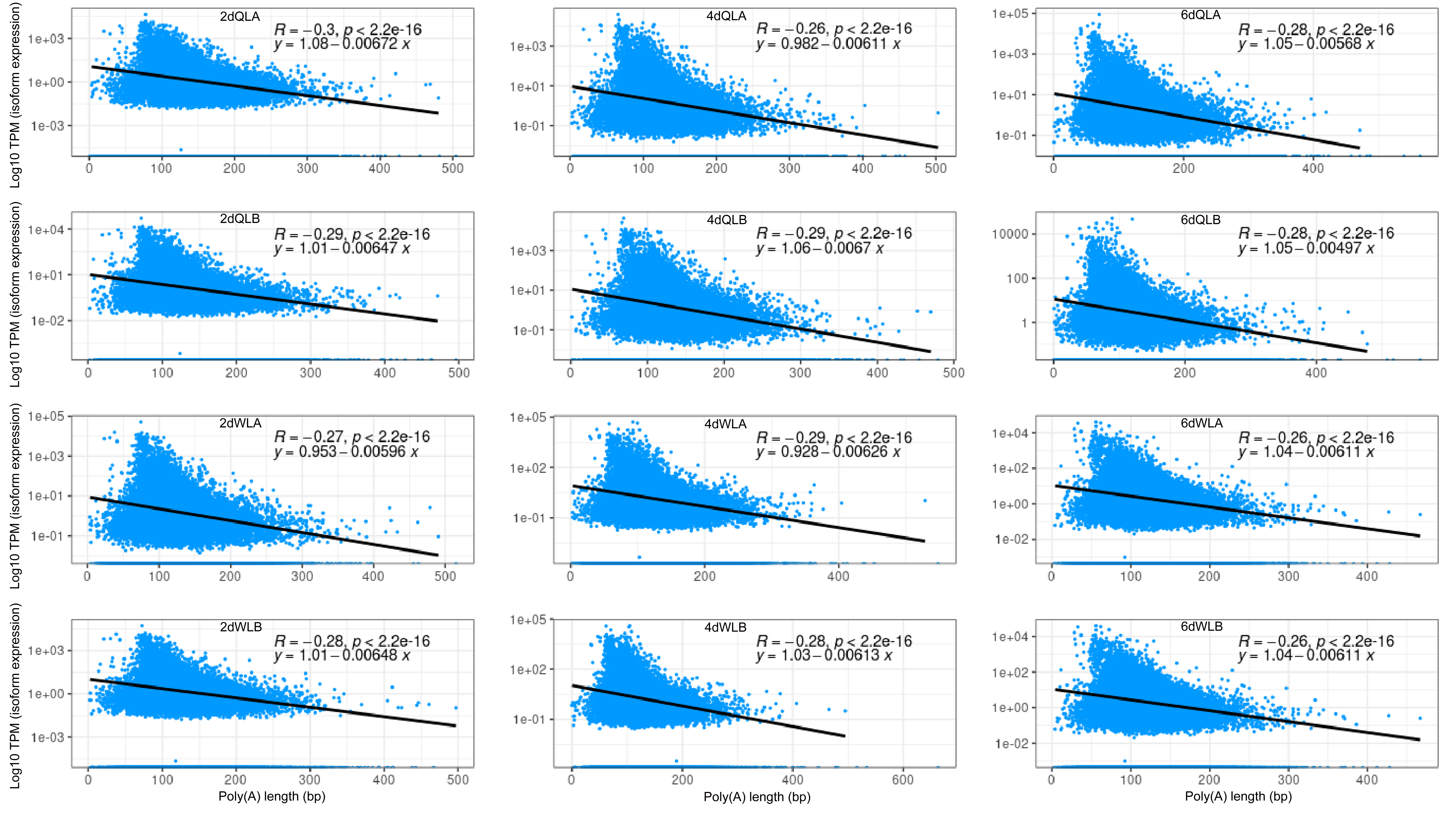

Fig.S5

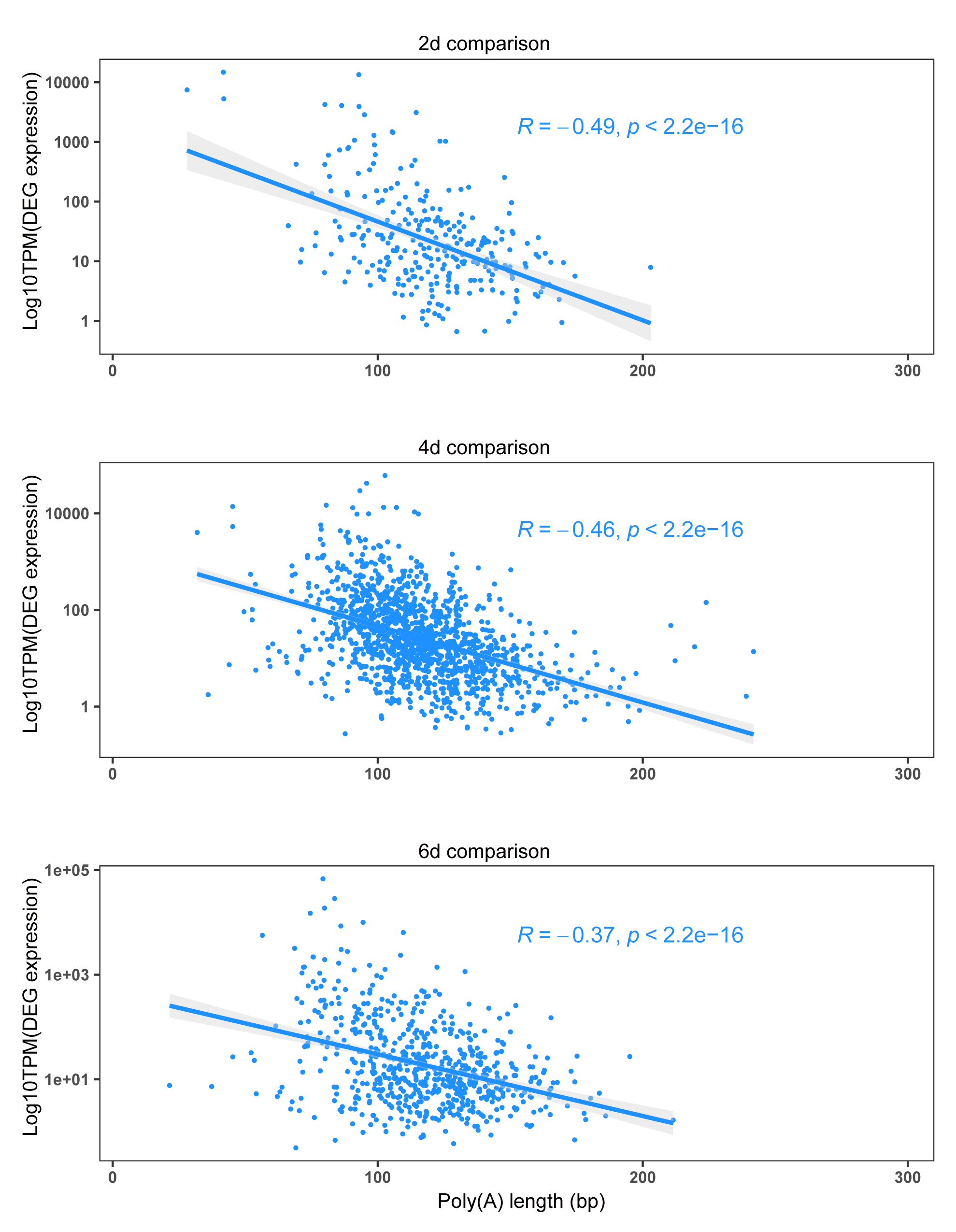

Fig.S6
